# Non-viral Intron Knockins Enable Simplified and Flexible Targeting of Endogenous Genes

**DOI:** 10.1101/2024.03.05.582227

**Authors:** Theodore L. Roth, Johnathan Lu, Alison McClellan, Oliver Takacsi-Nagy, Ansuman T. Satpathy

## Abstract

Targeting new genetic material to endogenous genes has opened diverse therapeutic and research applications, but current exon-based targeting methods have limited integration sites and are compatible only with complex or harsh selection methods. We present non-viral intron targeting, integrating large synthetic exons into endogenous introns to increase targeting flexibility and simplify selection of successfully edited cells. Engineered control of large synthetic exon’s splicing behavior further generalizes cell and gene therapy applications of non-viral intron knockins.

## MAIN TEXT

RNA guided nucleases such as Cas9 and Cas12a have dramatically expanded applications of endogenous gene editing. Combined with the addition of an exogenous DNA template, new genetic sequences can be added to defined sites in the genomes of primary human cells through homology directed repair (HDR). The ability to manipulate endogenous gene sequences and target new DNA to defined endogenous loci have opened diverse research and therapeutic applications, from directly correcting disease causing mutations, to integrating new synthetic genetic sequences with endogenous regulatory circuits such as by knockin of a Chimeric Antigen Receptor (CAR) or synthetic TCR to the endogenous TCR Alpha locus (TRAC) in primary human T cells^1,2^. Current gene targeting methods integrate new genetic material within exons of endogenous genes, normally knocking out the endogenous gene by targeting the middle of the gene’s coding sequence. Endogenous gene expression can be maintained by targeting the N or C terminus, although this limits gRNA target sites, resulting in effective editing in only a subset of potential target genes^3^.

Several methods exist to separate cells with a desired gene edit from unedited or incorrectly edited cells^4^. Classical selection methods such as positive selection using fluorescently conjugated antibodies and cell sorting (FACS), positive selection with antibodies conjugated to magnetic beads, or drug selection using resistance genes have recently been supplemented by selection strategies taking advantage of the ability to target transgenes to specific endogenous loci, such as by integration within essential genes^5–8^. However, each of these selection methods has significant drawbacks: (1) positive selections require direct cellular manipulation (such as antibody binding), which can leave bulky reagents bound to the cell surface after selection, inhibiting later cellular functions and presenting potential immunogenic antigens, (2) drug selections expose cells to toxic compounds and require expression of similarly potentially immunogenic foreign resistance genes, (3) both require additional genetic material beyond the desired therapeutic genetic sequence to be added to facilitate the selection, taking up large portions of the limited amounts of DNA deliverable into cells^9^, and (4) knockins to essential genes lose the benefits of choosing optimal endogenous gene regulation (e.g. CAR knockins to TRAC^1,2^). Touchless negative selections offer an ideal alternative, but for gene editing applications would require only successfully edited cells to lose expression of a selectable marker that all unedited and incorrectly edited cells retain.

We thus sought a generalizable method to negatively select successfully gene edited primary human cells. Traditional gene addition methods integrate new genetic material with an exogenous promoter to random (lenti/retrovirus) or non-expressed sites (safe harbors), or use gene editing to target an exon of an expressed gene so that the new sequence can be integrated into an existing mRNA under endogenous regulatory control. However, while these approaches are compatible with positive selection (by including additional DNA encoding a selection marker), they do not create the needed antigenic difference to enable negative selection. In the case of targeted editing of an essential gene’s exon, successfully HDR edited cells may lose expression of the targeted gene, but the majority of non-HDR edited cells will acquire NHEJ mutations which also results in target protein loss, as demonstrated by knockin of a CAR to the first exon of *TRAC* in primary human T cells (Fig. 1a)^1,2^.

**Figure 1.**
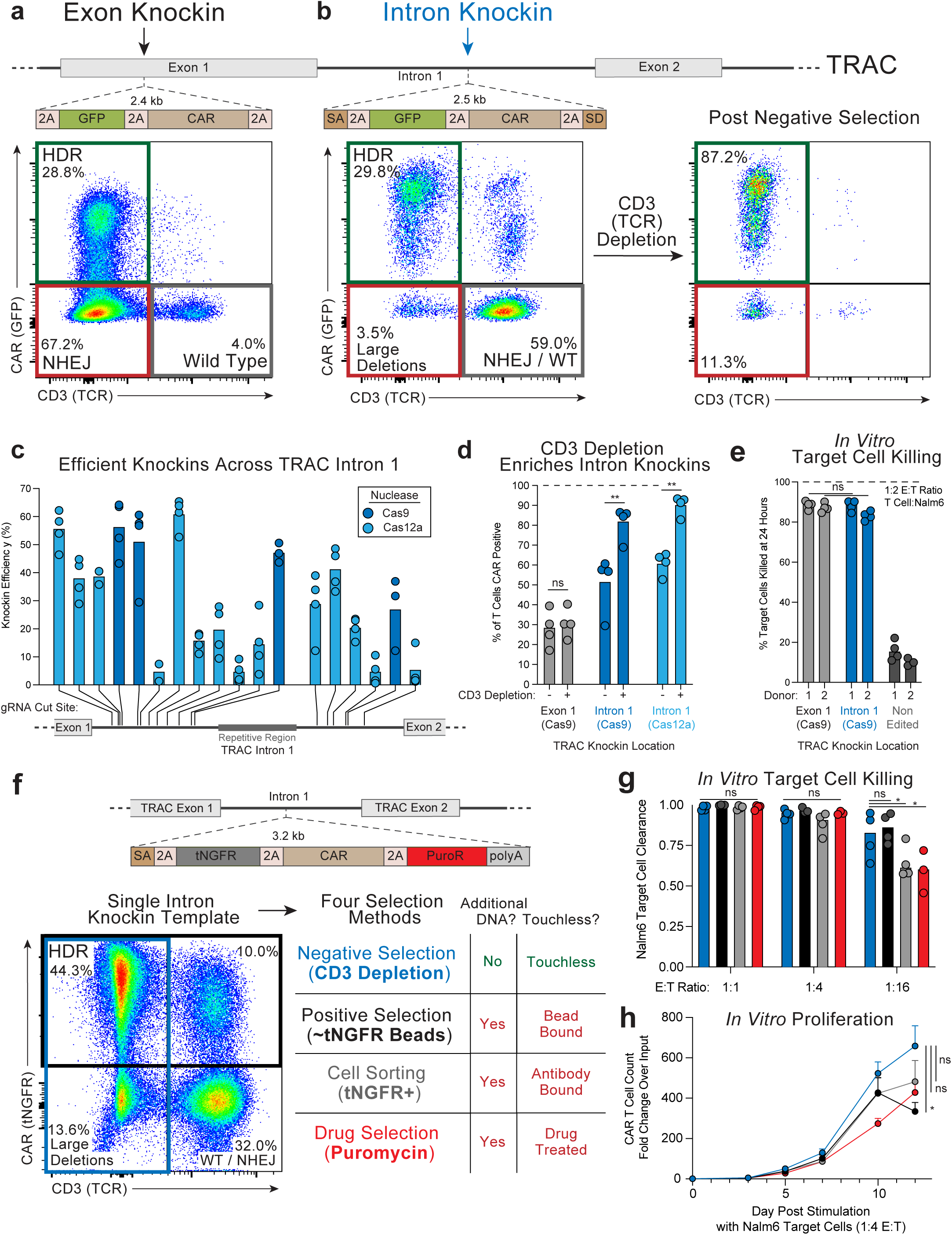
Non-viral TRAC Intron Knockins and Negative Selection of Successfully Edited CAR T Cells. **a,** Traditional endogenous gene targeting approaches integrate a new transgene into endogenous exons, causing knockout of the endogenous gene, as shown by knockin of a GFP-CAR multicistronic cassette into the first exon of *TRAC*, causing TCR loss in HDR edited cells. **b,** Knockin of a synthetic exon containing a GFP-CAR multicistronic cassette flanked by splice acceptor and donor sites within the first intron of *TRAC* shows the majority of non-HDR edited cells continue to express the targeted endogenous gene (TCR+ CAR-), while HDR edited cells have largely lost endogenous gene expression (TCR-CAR+). Successfully HDR edited cells can thus be negatively selected by depletion of TCR (CD3) expressing cells. **c,** Knockin efficiency of a synthetic exon containing a GFP-CAR multicistronic cassette across *TRAC* Intron 1 using both Cas9 and Cas12a nucleases. n = 2-4 unique donors per target site. **d,** CD3 depletion purifies successfully edited cells after Cas9 (>80% purity) and Cas12a (>90% purity) intron knockins, but provides only marginal enrichment for exon knockins due to undesired NHEJ edits causing TCR knockout. n = 4 unique donors. ns = not significant, ** = P<0.01, Paired t test. **e,** *TRAC* intron knockin CD19-28z CAR T cells kill Nalm6 target cells at same efficiency as *TRAC* exon knockin CAR T cells 48 hours post mixing at a 1:2 E:T ratio. Four technical replicates of n = 2 unique donors. ns = not significant, Mann-Whitney test. **f,** Intron knockin of a tNGFR-CAR-PuroR synthetic exon enabled a single population of edited cells to be compatible with four common selection methods. Only negative selections do not require additional genetic material (e.g. no selection marker or resistance gene necessary) and leave cells untouched post-selection. **g,** *In vitro* killing of Nalm6 target cells by *TRAC* intron knockin CD19-28z CAR T cells following negative, positive, sorting, or drug selection, measured 48 hours post co-incubation. n = 3-4 unique donors. ns = not significant, * = P<0.05, Paired t test. **h,** *In vitro* proliferation of *TRAC* intron knockin CD19-28z CAR T cells following negative, positive, sorting, or drug selection after co-culture with Nalm6 target cells. Error bars represent standard error of the mean from n = 3-4 unique donors. ns = not significant, * = P<0.05, Mann-Whitney test.

To overcome these issues with targeting flexibility and selection, we developed intron knockins, where synthetic exons are knocked into intronic regions of endogenous genes (Fig. 1b). Successful knockin to an endogenous intron results in splicing of the inserted sequence into the mature endogenous mRNA transcript, disrupting the endogenous gene and resulting in loss of endogenous protein expression (e.g. a surface protein for selection purposes), as used historically in gene-trap random mutagenesis screening methods^10,11^. But in contrast to exon targeting, NHEJ edits within introns are predominantly spliced out of mature mRNA transcripts^12^, resulting in continued protein expression (except for rarer large multi kb deletions^13^). Indeed, knockin of a CAR to the first intron of *TRAC* resulted in CAR positive, TCR negative cells, but unlike exon knockins, most CAR negative cells remained TCR positive (Fig. 1b). Negative selection of *TRAC* intron knockin CAR T cells by depletion of TCR (CD3) positive cells resulted in a dramatically enriched population of successfully edited CAR T cells.

Non-viral intron knockin of a CAR at the *TRAC* locus proved to be robust across intronic target sites, with efficient gene integrations possible using both Cas9 and Cas12a RNPs (Fig. 1c and Extended Data Fig. 1). Across unique human donors, *TRAC* intron knockins enabled negative selection of CAR T cells to greater than 90% purity (Fig. 1d). *TRAC* intron knockin CARs showed similar *in vitro* target cell killing as *TRAC* exon knockins (Fig. 1e). Finally, to directly compare different commonly used selection methods for primary human T cells, we designed a single synthetic exon that after TRAC intron knockin enabled the same population of bulk edited CAR T cells to be compatible with four distinct selection methods – touchless negative selection by anti-CD3 depletion, positive selection using magnetic beads, fluorescent cell sorting, and drug selection with puromycin (Fig. 1f and Extended Data Fig. 2). All four selection methods resulted in enriched and functional CAR T cell populations (Fig. 1g), with negatively selected CAR T cells showing a slight increase in *in vitro* proliferation after stimulation with target cells (Fig. 1h). Overall, TRAC intron knockins with negative selection enabled touchless purification of successfully edited CAR T cells without integration of any additional expressed exogenous DNA.

We next aimed to determine the extent to which intron knockins could accommodate integration of extremely large new genetic sequences. Past intron targeting approaches have focused on addition of short introns coding for small purification tags for biochemical experiments in cell lines^14,15^. Using a series of increasingly large synthetic exons, we found in primary human T cells that non-viral *TRAC* intron knockins were able to efficiently integrate functional synthetic exons of greater than 5 kb (Fig. 2a), which notably exceeds the packaging capacity of AAV vectors (Fig. 2b). Increasing sizes of intron knockins did show decreasing protein expression levels, although this was not specific to intron knockins as equivalently sized large knockins to *TRAC* exon 1 showed similarly lower levels of protein expression (Fig. 2c). Negative selection by CD3 depletion enabled enrichment of over 5kb intron knockins to greater than 50% purity on average (Fig. 2d), and *TRAC* intron edited CAR T cells bearing large synthetic exons maintained the same *in vitro* target cell killing capacity as *TRAC* exon targeted CAR T cells (Fig. 2e).

**Figure 2.**
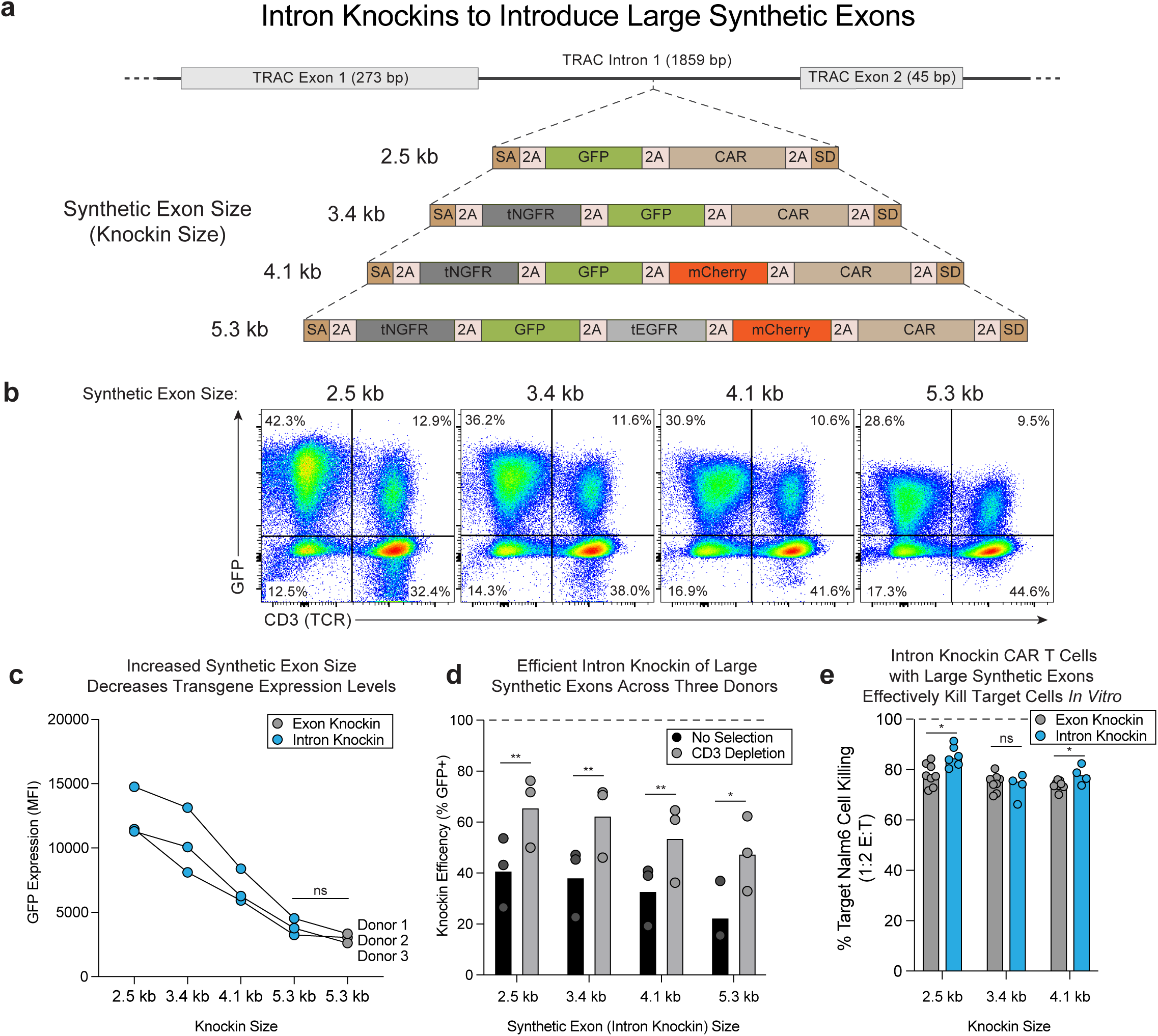
Introduction of >5 kb Large Synthetic Exons with TRAC Intron Knockins. **a,** Synthetic exons of increasingly large sizes from 2.5 kb to 5.3 kb targeted to *TRAC* intron 1. **b,** Flow cytometric plots demonstrating knockin efficiency of large synthetic exons at *TRAC* intron 1. **c,** Mean fluorescence intensity of GFP expression measured by flow cytometry in knockin positive cells after integration of increasingly large synthetic exons to *TRAC* intron 1. As the size of the synthetic exons increased, observed GFP expression decreased, potentially due to decreasing splicing efficiency of larger synthetic exons. n = 3 unique donors. ns = not significant, Paired t test. **d,** Knockin efficiency of increasingly large synthetic exons at *TRAC* intron 1 before and after negative selection by CD3 depletion. Even a 5.3 kb integration could successfully be enriched to greater than 50% knockin efficiency on average. n = 3 unique donors. * = P<0.05, ** = P<0.01, Paired t test. **e,** *TRAC* intron CAR T cells expressing a CD19-28Z CAR as part of increasingly large synthetic exons killed target Nalm6 tumor cells at equivalent efficiencies to *TRAC* exon CAR T cells with similar efficiencies to equivalent *TRAC* exon CAR T cells. Killing measured 48 hours after co-culture with a 1:2 E:T ratio. n = 2 unique human donors with 2-4 technical replicates each. ns = not significant, * = P<0.05, Unpaired t test. **a,** After intron knockin, successful splicing of a synthetic exon into the endogenous mRNA transcript disrupts the endogenous gene (“synthetic splicing”). If the synthetic exon is not spliced into the mature endogenous mRNA transcript, the new gene contained within the synthetic exon will not be expressed but endogenous gene expression is maintained (“endogenous splicing”). See Extended Data Fig 3a. **b,** Replacing the synthetic exon’s 3’ splice donor sequence with a polyA creates unbalanced splicing, resulting in the majority of cells expressing both the synthetic gene, e.g. GFP (“synthetic splicing”, Blue Box) while also expressing the targeted endogenous gene, e.g. TCR/CD3 (“endogenous splicing”, Orange Box). **c,** Varying the 3’ splicing architecture of synthetic exons allows different balances of synthetic splicing only (GFP+ TCR-) compared to synthetic and endogenous splicing (GFP+ TCR+). See Extended Data Fig 3b. n = 6 unique donors. ns = not significant, ** = P<0.01, Paired t test. **d,** Altering degenerate bases of the synthetic exon’s 5’ end (within the 2A multicistronic element necessary to separate the synthetic gene from the preceding endogenous exon’s translation) to include either Exonic Splice Silencer (ESS) or Exonic Splice Enhancer (ESE) elements similarly enables control over the degree of synthetic vs alternative splicing. **e,** Frequency of cells expressing both the synthetic knockin gene and the endogenous gene can be controlled by inclusion of ESS or ESE elements with the 5’ splicing architecture of synthetic exons. Note, the SV40 polyA construct in **b-c** contained the ESS element and its data is represented again for comparison with alternative 5’ splicing architectures. n = 2-6 unique donors. See Extended Data Fig. 3c. **f,** Intron knockins expand the toolbox for gene targeting methods. Intron knockins with 5’ and 3’ architectures that can alternatively splice between synthetic and endogenous exons allow more flexible targeting of synthetic genes under endogenous regulatory control, and intron knockins with splicing architectures resulting in only synthetic splicing enable touchless negative selection of successfully gene edited cells.

Finally, we reasoned that whereas exon knockins can only remove the endogenous gene (unless targeted to the N or C terminus, which often do not have suitable gRNA sites), intron knockins can be (1) alternatively spliced into a coding transcript, resulting in gene knockout, or (2) skipped, enabling the endogenous mRNA to be produced as well. We thus sought to determine whether alternative splicing of intron knockins could be synthetically controlled. By removing the 3’ splice donor site and replacing it with a polyA sequence, the splice sites present in the pre-mRNA become unbalanced, with the preceding endogenous exon’s splice donor able to splice with the integrated synthetic exon (“synthetic splicing”), or with the downstream endogenous exon (“endogenous splicing”) (Fig. 3a and Extended Data Fig. 3a). Indeed, while intron knockins with balanced splice sites resulted predominantly in loss of the targeted endogenous gene (GFP + / TCR -), intron knockins with unbalanced splice sites caused most knockin positive cells to express both the gene encoded in the synthetic exon, as well as the targeted endogenous gene (GFP+ / TCR+; Fig. 3b). The degree of alternative splicing between the synthetic and endogenous exons could be controlled by polyA choice, with the shorter SV40 polyA resulting in more endogenous splicing than the longer WPRE polyA, potentially due to greater ability of the longer polyA to halt transcription (Fig. 3c and Extended Data Fig. 3b). Rebalancing splicing through addition of a splice donor after the polyA sequence similarly decreased alternative splicing (Fig. 3c and Extended Data Fig. 3b).

**Figure 3.**
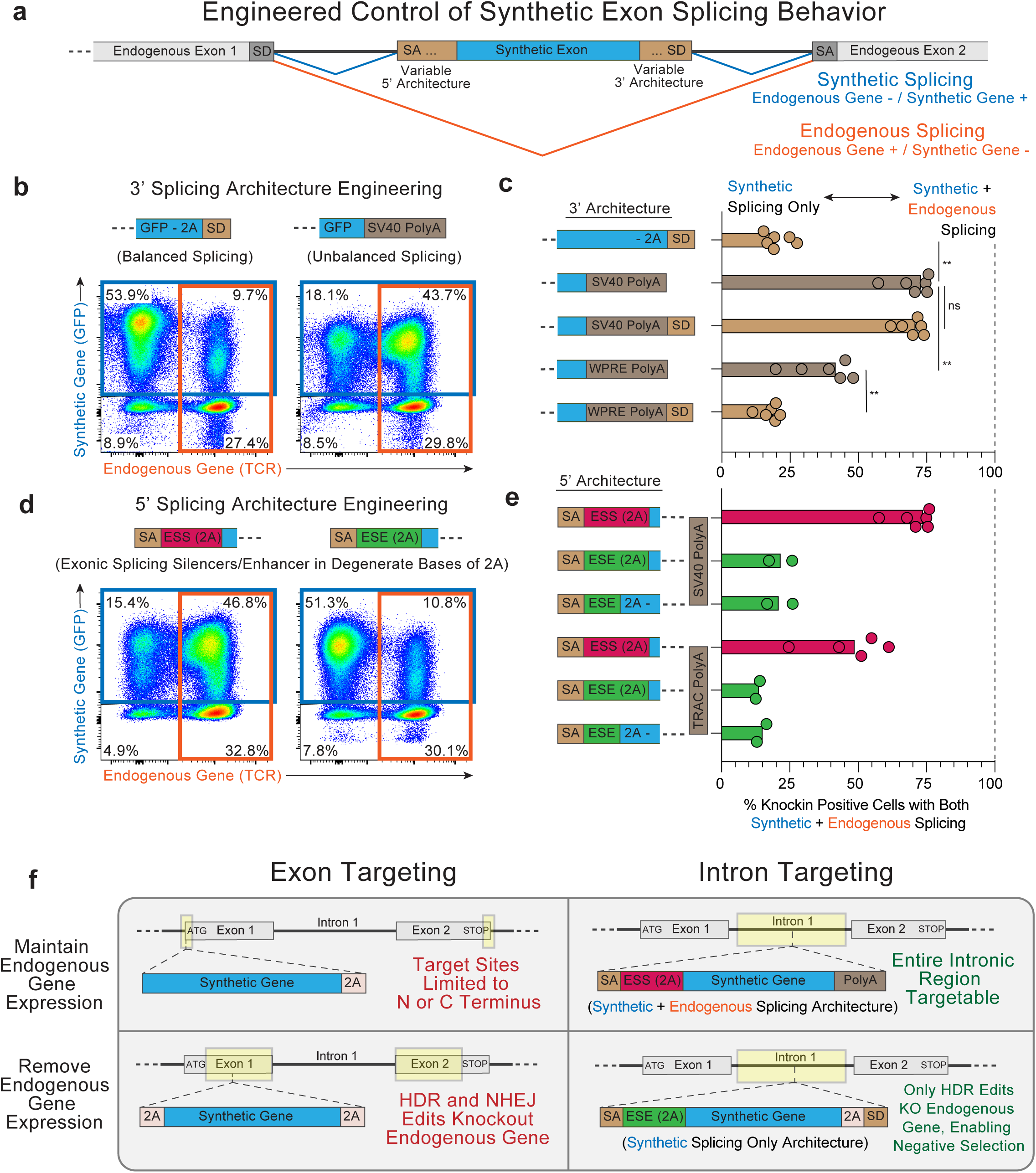
Engineered Control of Synthetic Exon Splicing using Intron Knockins. **a,** After intron knockin, successful splicing of a synthetic exon into the endogenous mRNA transcript disrupts the endogenous gene (“synthetic splicing”). If the synthetic exon is not spliced into the mature endogenous mRNA transcript, the new gene contained within the synthetic exon will not be expressed but endogenous gene expression is maintained (“endogenous splicing”). See Extended Data Fig 3a. **b,** Replacing the synthetic exon’s 3’ splice donor sequence with a polyA creates unbalanced splicing, resulting in the majority of cells expressing both the synthetic gene, e.g. GFP (“synthetic splicing”, Blue Box) while also expressing the targeted endogenous gene, e.g. TCR/CD3 (“endogenous splicing”, Orange Box). **c,** Varying the 3’ splicing architecture of synthetic exons allows different balances of synthetic splicing only (GFP+ TCR-) compared to synthetic and endogenous splicing (GFP+ TCR+). See Extended Data Fig 3b. n = 6 unique donors. ns = not significant, ** = P<0.01, Paired t test. **d,** Altering degenerate bases of the synthetic exon’s 5’ end (within the 2A multicistronic element necessary to separate the synthetic gene from the preceding endogenous exon’s translation) to include either Exonic Splice Silencer (ESS) or Exonic Splice Enhancer (ESE) elements similarly enables control over the degree of synthetic vs alternative splicing. **e,** Frequency of cells expressing both the synthetic knockin gene and the endogenous gene can be controlled by inclusion of ESS or ESE elements with the 5’ splicing architecture of synthetic exons. Note, the SV40 polyA construct in **b-c** contained the ESS element and its data is represented again for comparison with alternative 5’ splicing architectures. n = 2-6 unique donors. See Extended Data Fig. 3c. **f,** Intron knockins expand the toolbox for gene targeting methods. Intron knockins with 5’ and 3’ architectures that can alternatively splice between synthetic and endogenous exons allow more flexible targeting of synthetic genes under endogenous regulatory control, and intron knockins with splicing architectures resulting in only synthetic splicing enable touchless negative selection of successfully gene edited cells.

Alternative splicing could also be induced through manipulation of the 5’ end of the synthetic exon (Fig. 3d). Exonic Splicing Silencers (ESS) and Enhancers (ESE) are short ∼6-8 bp degenerate sequences that are bound by SR proteins to control the efficiency of mRNA splicing at adjacent splice acceptor sites^16,17^. We optimized the degenerate codons of the 2A multicistronic element at the 5’ end of the integrated synthetic exon (which separates the gene encoded by the synthetic exon from the preceding fragment of the targeted endogenous gene), systematically adding or removing predicted ESS and ESE sequences^18^. Across multiple constructs, the ESS-2A sequence induced alternative splicing, with both the synthetic and endogenous gene being expressed, whereas the optimized ESE-2A sequence predominantly resulted synthetic splicing only (Fig. 3e and Extended Data Fig. 3c). Biallelic intron knockin experiments further supported that a single targeted allele could generate both synthetically spliced and endogenously spliced mRNA transcripts (Extended Data Fig. 4).

Intron knockins expand the methodological toolbox for endogenous gene targeting in genetically modified cells for research and therapy (Fig. 3f and Extended Data Fig. 5). In addition to offering significantly more potential gRNA targets (∼30% of the genome is intronic compared to <3% coding), intron knockins also offer greater flexibility in the control of endogenous gene expression through alternative splicing. When expression of the targeted endogenous gene needs to be maintained, exon knockins can only target the N or C terminus, drastically limiting available gRNA options. In contrast, Intron knockins with engineered alternative splicing allow for target site selection across a gene’s entire intronic region. Moreover, when loss of the targeted endogenous gene is desired, intron knockins uniquely enable successfully edited cells to be purified by negative selection. Application of intron knockins in primary human T cells generated *TRAC* CAR knockin T cells that were touchlessly negatively selected to >90% purity without the need for additional bulky transgenes or requiring disruption of essential endogenous genes, and with complete removal of TCR positive cells. Indeed, similar findings for the ability to negatively select successfully edited CAR T cells at the *TRAC* locus using AAV-mediated intronic gene knockins were recently reported on bioRxiv, including the ability to perform multixplexed intron knockins at multiple endogenous surface receptors simultaneously^19^. Overall, Intron knockins offer a simple gene targeting strategy that overcomes key problems with prior selection methods for gene edited cellular therapies, while also dramatically expanding the flexibility of endogenous genetic manipulations.

## EXTENDED DATA FIGURE LEGENDS

**Extended Data Figure 1.**
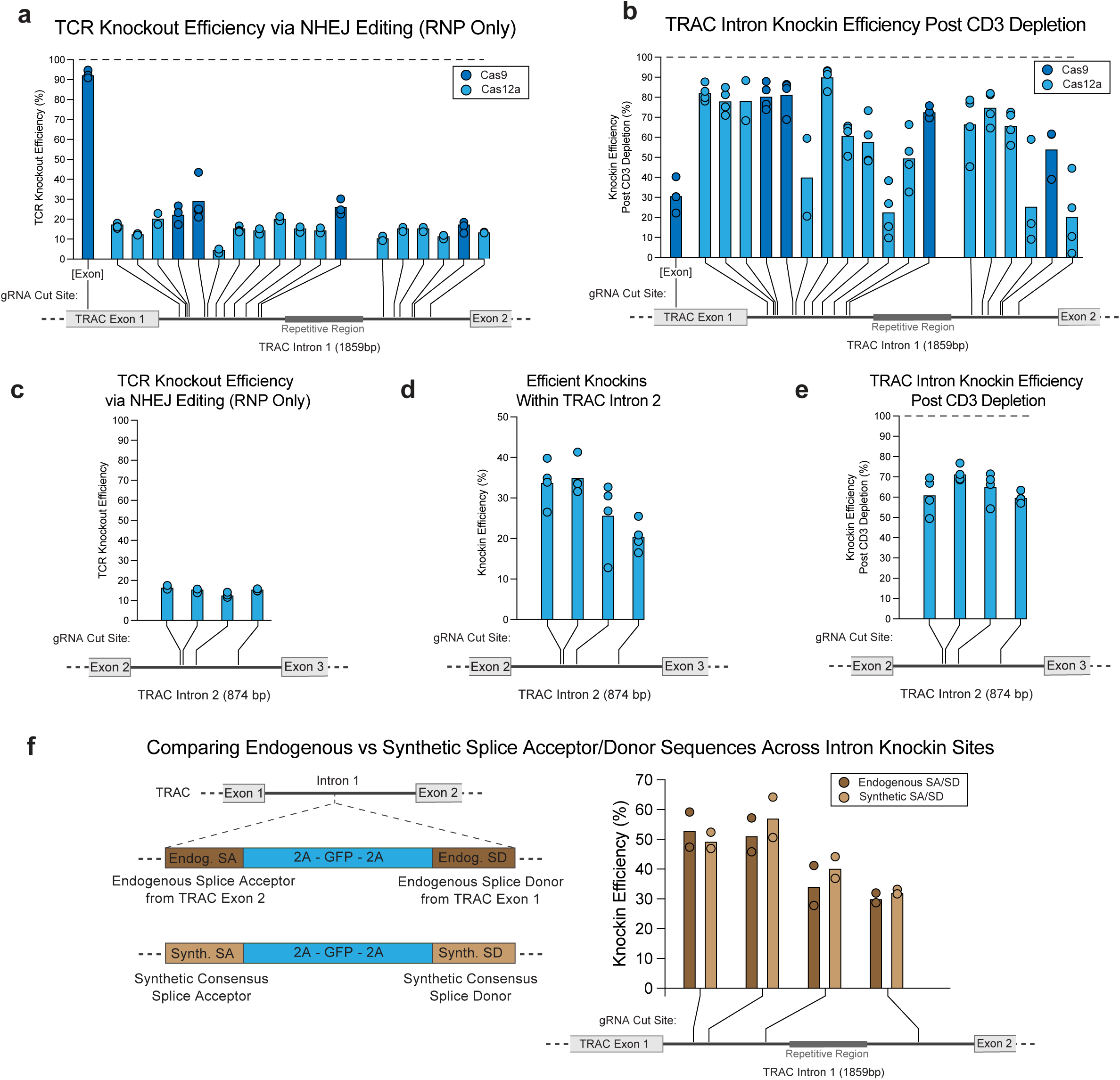
Efficient Knockin of a Synthetic Exon Across TRAC Intronic Sites. **a,** Observed efficiency of TCR knockout (measured by flow cytometric surface staining for CD3) after gene editing with Cas9 or Cas12a RNPs containing gRNAs targeting *TRAC* exon 1^2^ or 18 distinct targets within *TRAC* intron 1 (avoiding a highly repetitive region within the intron). TCR knockout was highly efficient for the exon targeting gRNA, but significantly lower for intronic guides. **b,** Percentage of total cells expressing the knockin gene cassette (GFP+) following CD3 depletion (removing TCR positive cells) after intron knockin at 18 unique sites throughout *TRAC* intron 1. n = 2-4 unique donors. **c,** TCR knockout (measured by flow cytometric surface staining for CD3) after gene editing with Cas12a RNPs containing gRNAs targeting *TRAC* intron 2. **d,** Knockin efficiency at four unique sites within TRAC intron 2. **e,** Percentage of total cells expressing the knockin gene cassette (GFP+) following CD3 depletion (removing TCR positive cells) after intron targeting at 4 unique sites throughout *TRAC* intron 2. **f,** Across four *TRAC* intron 1 target sites targeted with Cas9 RNPs, no major difference in knockin efficiency and successful protein expression from the integrated synthetic exon was observed when using the endogenous splice acceptor and donor sites from the adjacent endogenous *TRAC* exons compared to synthetic consensus splice acceptor and donor sites. **a-f,** n = 2-4 unique primary human T cell donors.

**Extended Data Figure 2.**
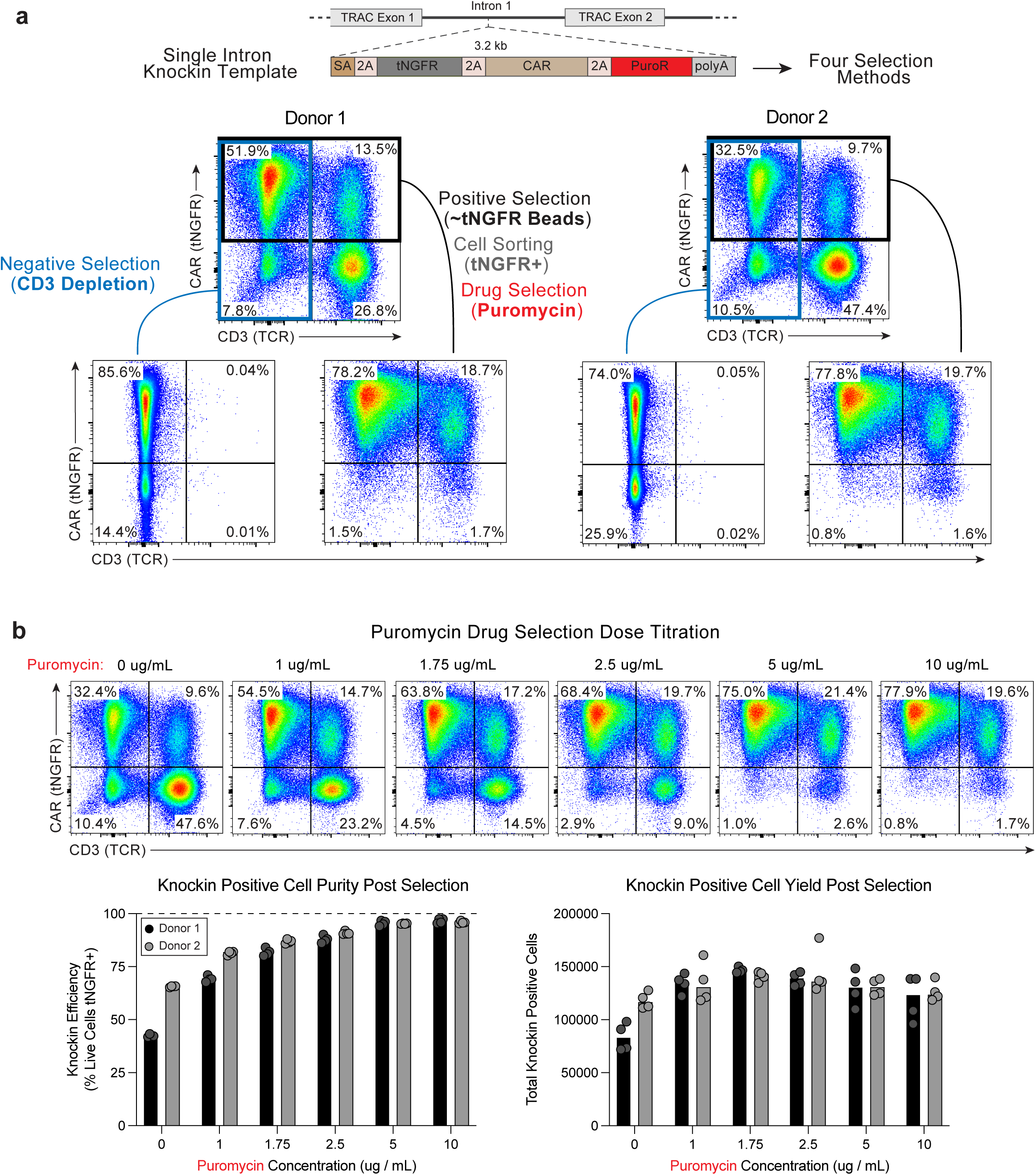
Direct Comparison of Diverse Selection Methods using TRAC Intron Knockins. **a,** Representative flow cytometric plots of TRAC intron CAR T cells after purification with four distinct selection methods. Knockin of a synthetic exon containing a tNGFR-CAR-PuroR multicistronic cassette to *TRAC* intron 1 enabled successfully edited cells to be negatively selected by CD3 Depletion (removal of TCR positive cells), positively selected by streptavidin magnetic bead enrichment after binding of anti-tNGFR biotinylated antibodies, fluorescence-activated cell sorting after binding of anti-tNGFR fluorescent antibodies, or drug selection after culture in puromycin. Negative selection by CD3 depletion yields predominantly a successfully edited CAR+ T cell population without the endogenous TCR, although rarer TCR negative / knockin negative cells are present (likely due to the RNP induced double stranded break within the intron causing a large deletion that included a portion of one or both adjacent exons, instead of the more common NHEJ repair outcome of smaller indels). Positive selection, sorting, and drug selection in contrast remove all knockin negative cells, but retain a population of TCR positive / knockin positive cells (likely due to successful HDR mediated knockin to one TRAC allele with either no editing or a small indel removed during mRNA splicing on the second TRAC allele; while in some T cells one TCRα loci is silenced, numerous T cells express functional TCRα chains from both alleles^20^). **b,** Puromycin dose titrations reveal a tradeoff between purity and cell yield when performing drug selections. Increasing doses of puromycin yielded greater T cell purity, but overall edited cell yield began to decline with increasing puromycin concentrations. Beginning 6 days post editing, 100,000 bulk edited T cells were treated with indicated doses of puromycin for 48 hours. A dose of 5 ug/mL was used for experiments in Fig. 1f-g to balance purity and yield. n = 2 donors with 4 technical replicates each.

**Extended Data Figure 3.**
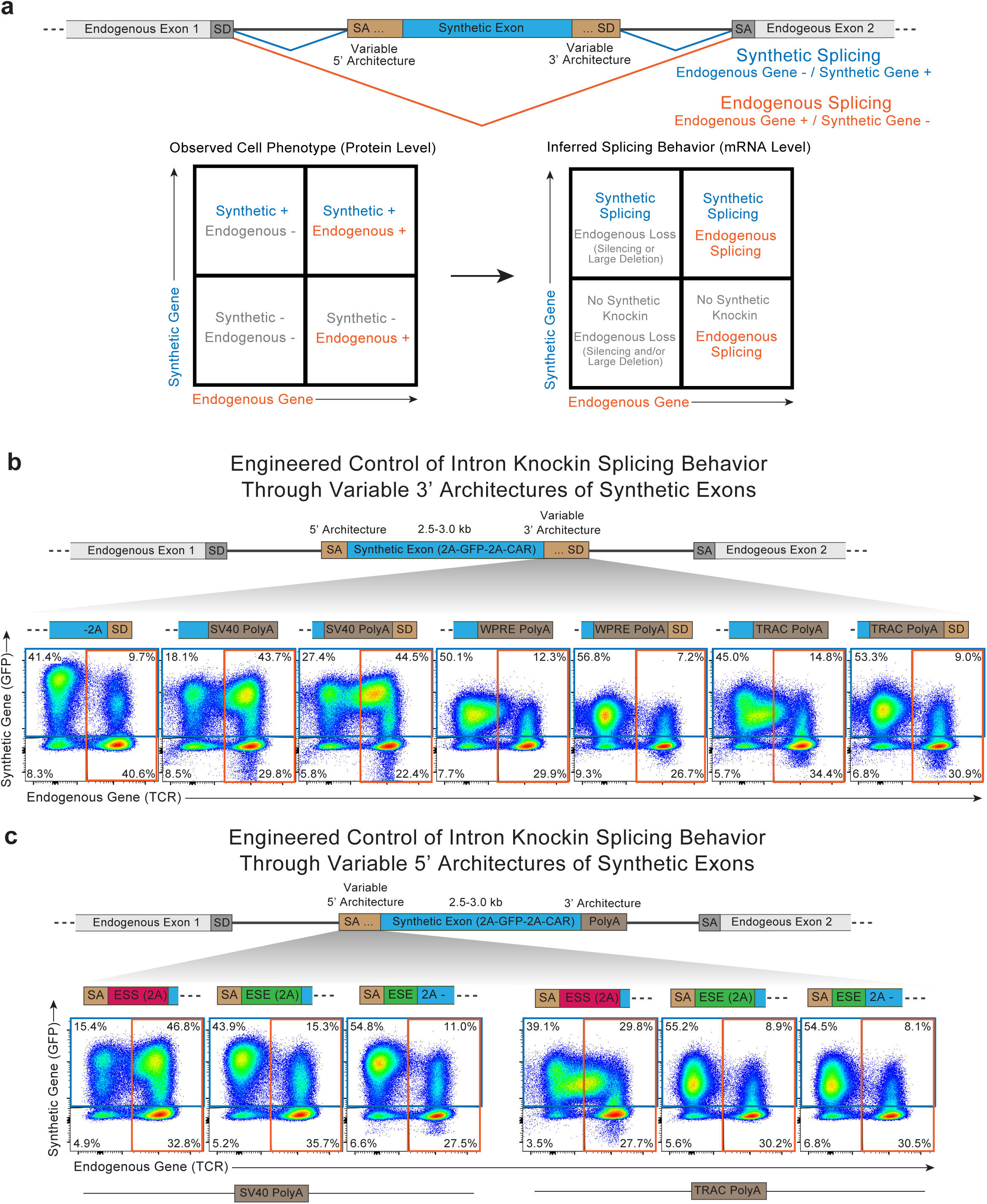
Variable 5’ and 3’ Splicing Architectures Enable Control of Alternative Splicing of Synthetic Exons. **a,** Correlation between observed cellular protein phenotypes by flow cytometry with inferred splicing behavior. Cells expressing both the protein encoded by the synthetic exon (e.g. GFP) and the endogenous protein (e.g. TCR) must be undergoing both synthetic splicing and endogenous splicing. These two splicing outcomes could be occurring from the same pre-mRNA transcript from a single allele by alternative splicing (single pre-mRNA transcript spliced into two different mature mRNA transcripts, one encoding the synthetic gene and a second encoding the endogenous gene), or the two splicing outcomes could be occurring within two separate pre-mRNA transcripts expressed from two different alleles (e.g. one allele with successful intron knockin of the synthetic exon, with the second allele being either unedited or possessing small indels from NHEJ repair that do not interfere with endogenous gene expression). Biallelic intron knockin experiments (Extended Data Fig. 4) support that the majority, but not all, dual positive cells result from alternative splicing of the same pre-mRNA from a single allele. **b,** Variable 3’ synthetic exon splicing architectures lead to controllable degrees of alternative splicing. Balanced splicing (2A-SD, far left) results in the majority of knockin (GFP) cells being negative for the endogenous gene (TCR-), with the residual GFP+ TCR+ cells likely due to one knockin allele and one WT or NHEJ edited allele. Creation of unbalanced splicing by replacement of the splice donor with a polyA (either SV40, WPRE, or *TRAC’s* endogenous polyA^21^) resulted in an increase in dual positive cells, with varying levels of endogenous gene expression depending on the polyA sequence used (SV40 > WPRE or *TRAC*). The differing levels of endogenous gene expression observed were potentially due to variable efficiency of RNA transcription termination by the polyA sequences, with the shorter SV40 polyA sequence less efficient at stopping transcription, enabling more mRNA transcripts to continue to the endogenous exons downstream from the knocked in synthetic exon and thus be capable of alternative splicing. Indeed, addition of a splice donor sequence following the polyA tails decreased the expression levels of the endogenous gene for all three polyA sequences, returning completely to baseline (2A-SD construct, far left) with the WPRE and TRAC polyA, and lowering the TCR expression level for the SV40 polyA, supporting the interpretation that unbalanced splice sites in the polyA only constructs lead to greater alternative splicing. SV40 polyA flow plot reproduced from Fig. 3b for comparisons. **c,** Alternative splicing can also be controlled by varying the synthetic exon’s 5’ splicing architecture. In intron knockin constructs containing both an SV40 or *TRAC* endogenous polyA sequences at their 3’ end, inclusion of Exonic Splicing Silencer (ESS) DNA sequences adjacent to the synthetic exon’s 5’ splice acceptor resulted in increased numbers of dual positive cells showing evidence of alternative splicing. The short 6-8 bp ESS sequences were introduced into the 2A multicistronic element at the 5’ end of the synthetic exon (necessary to separate the new synthetic gene’s protein translation from the translation of the preceding endogenous exon) using degenerate bases to maximize the number and strength of the ESS sequences present^18^. In pre-mRNA transcripts containing unbalanced splice sites due to the 3’ polyA, inclusion of ESS sequences at the synthetic exon’s 5’ end reduces binding by the SR proteins that mediate splicing and increases the chance that the preceding endogenous exon will splice with the downstream endogenous exon rather than the synthetic exon. Both splicing outcomes occur with alternative splicing, resulting in expression of both the endogenous gene and the new synthetic gene. In contrast, similar optimization of the 2A elements degenerate bases to include Exonic Splicing Enhancer elements (or insertion of in-frame ESE elements prior to the unaltered 2A sequence) largely prevented alternative splicing, returning the levels of dual positive cells back to amount seen with synthetic exons containing balanced splice sites (e.g. the amount of dual positive cells seen due to one knockin allele and one unedited/NHEJ allele). ESS-2A – SV40 polyA flow plot reproduced from Fig. 3d for comparisons.

**Extended Data Figure 4.**
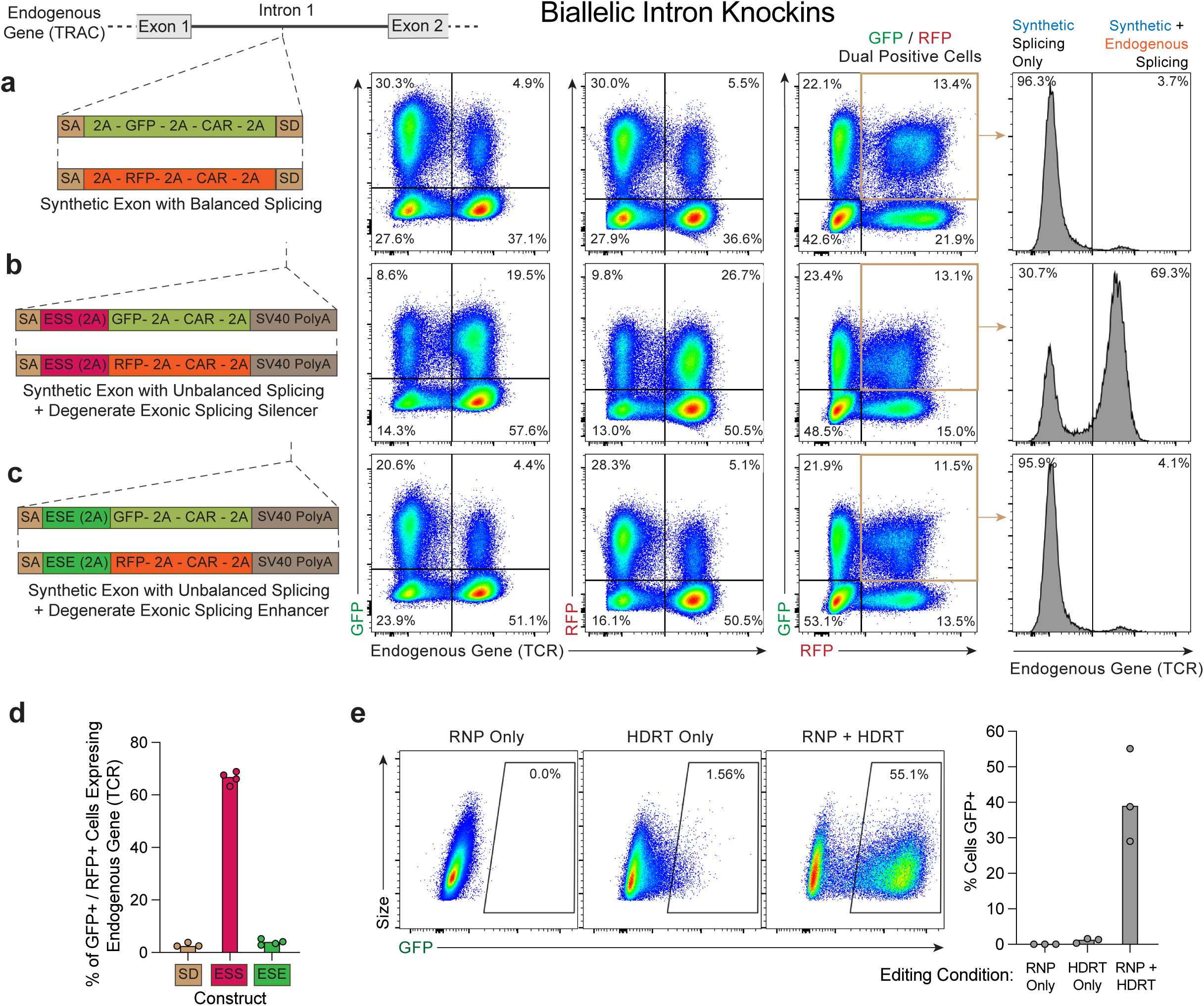
Biallelic TRAC Intron Knockins Demonstrate Alternative Splicing of Synthetic Exons. **a,** Simultaneous intron knockin of two synthetic exons containing two separate fluorescent proteins (GFP and RFP/mCherry) enables dual positive cells with biallelic edits to be identified. With knockin of synthetic exons with balanced splice sites, gating on dual positive cells showed almost none of the dual GFP positive / RFP positive cells still expressed the endogenous gene (TCR, measured by CD3 staining). **b,** Biallelic knockin of a synthetic exon containing Exonic Splicing Silencing elements at its 5’ end and a polyA at its 3’ end (creating unbalanced splice sites) showed dual GFP positive / RFP positive cells to be largely TCR positive. As both TRAC alleles possess a knocked in synthetic exon in the dual positive cells, one or both alleles must be capable of alternative splicing to also still maintain expression of the endogenous gene (e.g. to be TCR positive). **c,** Biallelic knockin of a synthetic exon containing Exonic Splicing Enhancer elements at its 5’ end do not show evidence of alternative splicing, with almost all dual GFP positive / RFP positive cells negative for the endogenous gene. The ability of a handful of silent degenerate basepair changes at the 5’ end of the synthetic exon (ESS → ESE element) to drastically change the degree of dual positive cells expressing the endogenous gene (TCR) further supports that a single allele with an intron knockin is capable of either expressing only the gene encoded in the synthetic exon (using splicing architectures in **a** and **c**), or is capable of alternative splicing and expression of both the synthetic gene and the endogenous gene (using splicing architecture in **b**). **d,** Quantification of the percentage of dual GFP positive / RFP positive cells that continue to express the endogenous gene across the three tested splicing architectures in **a-c**. n = 3-4 unique donors. **e,** The degree of off-target integrations is important for interpreting biallelic integration data, as dual positive cells also expressing the endogenous gene (GFP / RFP / TCR positive cells) could be due to one knockin allele, one unedited endogenous allele, and then an off-target integration of the second fluorescent protein. The degree of off-target vs on-target integrations was assessed by electroporation of the GFP DNA Homology Directed Repair Template used in **b** without its accompanying RNP. While a low amount of GFP expression was seen in ∼1-2% cells with HDRT only electroporations, likely due to off-target integrations, this was markedly less than the knockin rate and expression level of GFP when the HDRT was electroporated with its on-target RNP, similar to prior studies^2^. The low frequency of off-target integrations could not account for the degree of endogenous gene (TCR) positivity in the dual GFP / RFP positive cells in **b**. n = 3 unique donors.

**Extended Data Figure 5.**
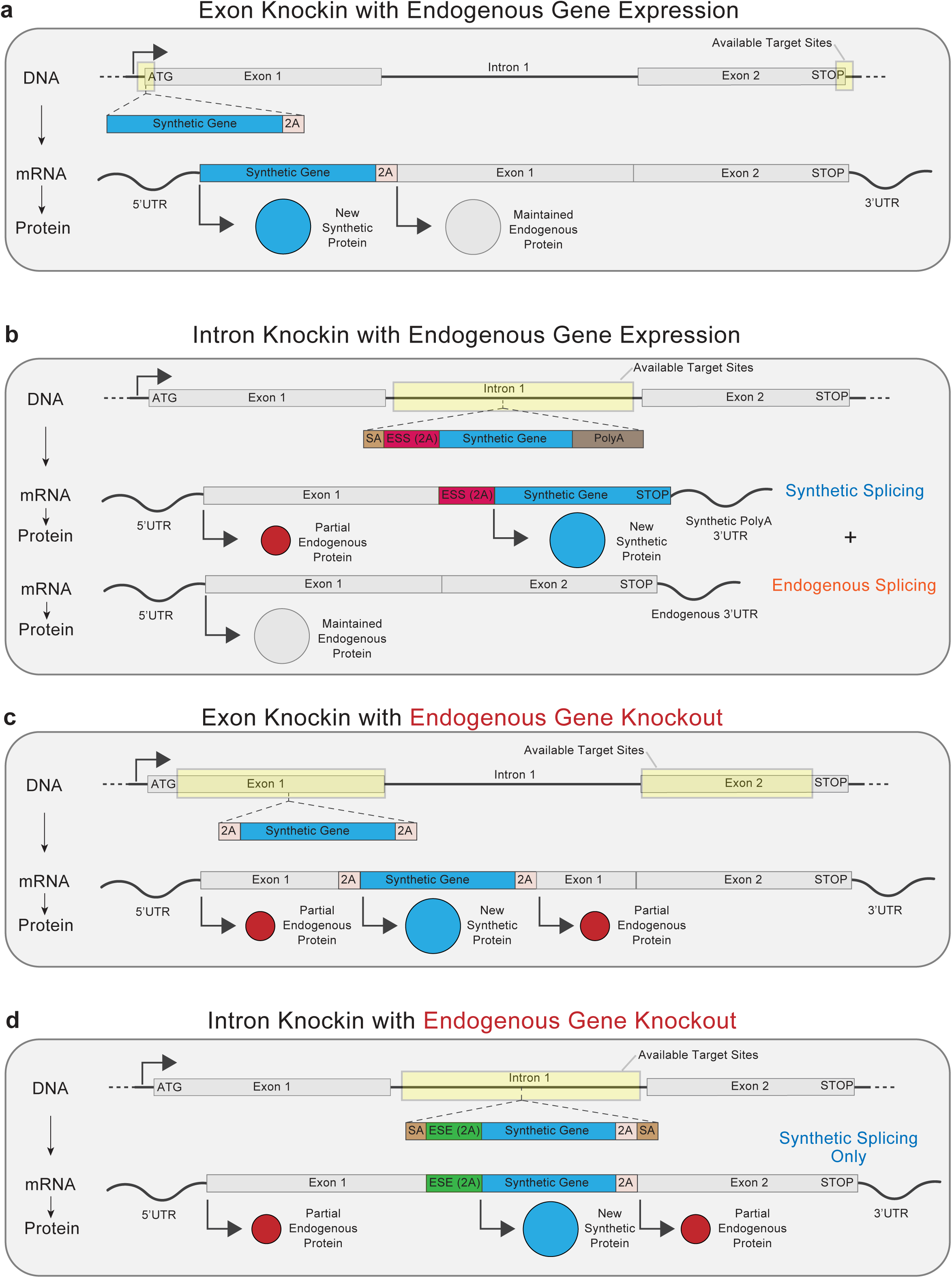
Generalized Endogenous Gene Targeting with Exon and Intron Knockins. **a,** A new synthetic gene can be introduced under endogenous regulatory control of an existing gene without also loosing expression of the existing gene only be integration at the N terminus (immediately before the Start codon) or C terminus (immediately before the Stop codon) of the targeted gene. This limitation in target sites means that efficient gRNAs for gene knockin may not be present for many genes, as observed in previous studies^3^. **b,** Intron knockins offer greater flexibility for placing a synthetic gene under endogenous regulatory control without disrupting the endogenous gene. Using optimized 5’ and 3’ splicing architectures to induce alternative splicing, a synthetic exon can be introduced throughout the intronic regions of the endogenous gene, with alternative splicing resulting in two separate mature mRNA transcripts, one encoding the new synthetic protein and a second encoding the endogenous gene. Orders of magnitude more gRNA target sites are available within intronic regions than only at the very N or C terminus of a gene. **c,** Exonic targeting of a new synthetic gene within the coding sequence of an endogenous gene disrupts the endogenous gene’s sequence, resulting in a multicistronic mRNA transcript encoding the new synthetic protein and two partial fragments of the endogenous protein, causing knockout of the endogenous gene. **d,** Intron targeting with a synthetic exon containing optimized 5’ and 3’ splicing architectures to induce synthetic splicing only also results in endogenous gene knockout. However unlike in exonic targeting, where alleles with unsuccessful knockins largely have NHEJ induced indels resulting in frameshift mutations and protein knockout, with intron targeting NHEJ induced indels reside within an intronic sequence that is largely tolerant of short DNA changes which will be spliced out of the final mRNA transcript (more rare but detectable larger deletions that include parts of the surrounding exons induced by double stranded breaks can still result in endogenous protein knockout). If the targeted endogenous gene is a surface receptor, then intron knockins offer the unique ability to perform negative selections to purify successfully edited cells.

## METHODS

### Primary Human T Cell Isolation and Culture

PBMCs from healthy human blood donors were collected under an approved IRB protocol by the Stanford Blood Center and used to isolate human T cells. Briefly, leukoreduction chambers (LRS) from processing of platelet donations were used to isolate PBMCs using density centrifugation with Ficoll (Lymphoprep, StemCell) within SepMate tubes (StemCell) according to manufacturers instructions. Next, primary human CD3 positive T cells were isolated by negative selection using Human CD3 T Cell Enrichment kit (StemCell) according to manufacturer’s instructions. Isolated primary human CD3 T cells were counted using an automated cell counter (Countess, Thermo), and activated using anti-human CD3/CD28 dynabeads (Cell Therapy Systems, Thermo) at a 1:1 ratio in XVivo 15 media (Lonza) supplemented with 5% FBS (MilliporeSigma) and 50 U/mL of human IL-2 (Peprotech). T cells were activated at 1:1 ratio of cells to dynabeads, and initially cultured in standard tissue culture incubators at approximately 1e6 cells / mL media. After gene editing/electroporations, T cells were counted and reseeded at approximately 1e6 cells / mL XVivo 15 media with fresh IL-2 every 2-3 days.

### Non-viral gene knockins

Two days after activation, human T cells were harvested, dynabeads were magnetically removed by incubating for two minutes at room temperature on a magnet (EasySep Magnet, StemCell), and cells were counted using an automated cytometer. For electroporations, one million T cells per editing condition were gently pelleted by centrifugation at 90G for 10 minutes, followed by careful aspiration of the supernatant. T cell pellets were resuspended in 20 uL per editing condition in P3 Buffer (Lonza) and then mixed with prepared RNP and DNA HDRT templates. For each Cas9 knockin condition, RNPs were prepared by first complexing the gRNA by mixing 0.375 uL of 200 uM tracrRNA (IDT) with 0.375 uL of 200 uM crRNA (IDT) and incubating for 15 minutes at room temperature. Next 0.25 uL of 100 mg/mL PGA (15–50 kDa poly(L-glutamic acid); MilliporeSigma) was then added to the complexed gRNA and mixed by pipetting up and down. Next 0.5 uL of 40 uM SpCas9 (UC Berkeley MacroLab) was then added, mixed by pipetting up and down, and incubated for 15 minutes at room temperature to form the final Cas9 RNP. For Cas9 knockins, 20 uL of T cells were mixed with 1.5 uL of RNP (60 pmols total RNP) and 4 uL of plasmid DNA HDR Template at 1 ug/uL (4 ugs total HDRT). For each Cas12a condition, RNPs were prepared by first mixing 0.4 uL of 200 uM Cas12a gRNA (IDT) with 0.2 uL of 100 mg/mL PGA and pipetting up and down. Next 0.4 uL of 60 uM AsUltraCas12a (UC Berkeley MacroLab) was added and mixed by pipetting up and down, followed by incubation at room temperature for 10 minutes. For Cas12 a knockins, 20 uL of T cells were mixed with 1 uL of Cas12a RNP (24 pmols total RNP) and 4 uL of plasmid DNA HDR Template at 1 ug/uL (4 ugs total HDRT). For both Cas9 and Cas12a knockins, T cells were electropoated on a Gen2 Lonza 4D electroporation/nucleofection system using 96 well plate attachment and 20 uL cuvettes, using pulse code EO-151. Immediately following electroporation, 75 uL of pre-warmed XVivo 15 media was added to each cuvette, and cells were rested within the cuvettes for 15 minutes in a standard 37C Tissue Culture Incubator prior to moving to culture plates or flasks. An annotated list of all gRNA sequences used in the study is available in Supplementary Table 1.

### DNA Constructs and HDR Templates

All DNA constructs used in the study were generated using standard cloning methods, primarily using PCRs (Q5 ultra II HotStart Polymerase, NEB), Gibson Assemblies (NEBuilder HiFi DNA Assembly Master Mix, NEB) and bacterial transformations (Mix & Go! Competent Cells - DH5 Alpha, Zymo). DNA constructs were sequence confirmed by Sanger Sequencing (Elim Bio) or whole plasmid next-generation sequencing (Primordium). For gene editing, plasmid Homology Directed Repair Templates were produced by bacterial culture and standard plasmid preparation (Zymo Midiprep and Maxiprep kits) according to manufacturer instructions, including endotoxin removal steps. Final plasmid HDR templates were eluted in molecular grade water, quantified (Nanodrop), and diluted to final concentrations of 1 ug/uL for electroporations. An annotated list of all DNA constructs used in the study, as well as full DNA sequences for all constructs, is available in Supplementary Table 2. Annotated genebank files for all DNA constructs used in the study are available upon request.

### Flow Cytometry

All flow cytometric data shown was acquired either within the Stanford Blood Center Flow Cytometry Core using a BD Fortessa analyzer, or using a Biorad ZE5 analyzer. All antibodies used for flow cytometric experiments and staining dilutions are listed in Supplementary Table 3. Briefly, for flow cytometric staining, samples of approximately 100,000 T cells per analyzed condition were placed into round bottom 96 well plates and centrifuged for 5 minutes at 300G. After discarding the supernatant, each well was resuspended in 20 uL of FACS Buffer (PBS + 2% FBS) mixed with desired antibodies at indicated dilutions (Supplementary Table 3), and incubated at 4C for 20 minutes in the dark. Cells were washed once with FACS Buffer and resuspended in FACS Buffer for analysis. Flow cytometric data was analyzed using FlowJo v10 software.

### Negative Selection by CD3 Depletion

Negative selections for successfully intron edited T cells were performed by CD3 depletion. Briefly, six days after non-viral gene editing by electroporation, subsequently expanded T cells were harvested, counted, and anti-CD3 biotinylated antibodies were added, following manufacturer’s instructions (EasySep™ Human CD3 Positive Selection Kit II, StemCell). After addition of magnetic beads and incubation at room temperature on a magnet, the supernatant containing untouched, CD3 negative cells was poured off into a new tube. Magnetic beads were added a second time according to manufacturer’s instructions, and again the supernatant was poured off after incubation on a magnet at room temperature, resulting in a final population of negatively selected, untouched, CD3 negative T cells.

### Positive Selection by ∼tNGFR Magnetic Bead Purifications

Positive selection for successfully intron edited T cells were performed by antibody binding of the surface expressed selection marker tNGFR followed by conjugation to a magnetic bead for magnetic purification six days after non-viral gene editing. Briefly, an anti-tNGFR biotinylated antibody (Supplementary Table 3) was bound to the surface of T cells using the same staining procedure as described above for flow cytometric fluorescent antibody staining. After antibody binding, T cells were incubated with magnetic beads coated with streptavidin (Dynabeads MyOne Streptavidin T1, Thermo) at 1.25 uL of beads per 1e6 cells. T cells were then incubated at room temperature on a rotating mixture for 15 minutes. Bound T cells were then placed onto a magnet and incubated for 5 minutes before pouring off the supernatant. T cells were resuspended in FACS Buffer and placed on the magnet a second time. After pouring of the supernatant, T cells were resuspended in XVivo 15 culture media containing 1 uM free biotin (D-Biotin, Thermo) for 2 days to competitively compete for streptavidin binding sites and help to remove bound magnetic beads from the T cell surface before returning cells to standard culture conditions.

### Fluorescent Cell Sorting

Sorting based selection of successfully intron edited T cells was performed using standard fluorescence activated cell sorting six days after non-viral gene editing. Briefly, cells were bound with an anti-tNGFR fluorescent antibody (Supplementary Table 3) for sorting as described for flow cytometric staining. A BD Aria Cell Sorter (Stanford Blood Center Flow Cytometry Core) was used for all cell sorting experiments. Cells were maintained at 4C throughout the duration of sorting, and sorted cells were collected into destination tubes of XVivo15 media mixed 1:1 with FBS. After sorting, cells were centrifuged for 5 min at 300G prior to resuspension in culture media.

### Drug Selection with Puromycin

Drug based selection of successfully intron edited T cells was performed using puromycin selection for forty-eight hours beginning six days after non-viral gene editing. Briefly, T cells were cultured in XVivo15 based media as described above with the addition of 5 ug/mL of puromycin (Puromycin Dihydrochloride, stock concentration of 10 mg/mL in 20 mM HEPES buffer, ThermoFisher). Concentrations used in dose titration experiments ranged from 1.0 ug/mL to 10 ug/mL as indicated in Extended Data Fig. 2b. After two days of puromycin selection, cells were centrifuged for 5 minutes at 300g and the supernatant was removed before resuspension in standard culture media without puromycin.

### In Vitro Cancer Target Cell Killing Assays

At eight days post non-viral gene editing, edited CAR T cells (either bulk populations without selection or selected cells as described) were mixed at indicated E:T ratios with target Nalm6 leukemia cells in 96 well plates, with 40,000 Nalm6 cells and varying numbers of T cells per well. Cell killing was assessed by flow cytometery at 48 hours, and the percentage of Nalm6 tumor cell killing was calculated by taking 1 – (# Nalm6 cells alive in experimental condition / # Nalm6 cells alive in no T cell conditions). Nalm6 leukemia cells (ATCC) were cultured in RPMI+10% FBS and passaged every 2-3 days to maintain cell densities of approximately 1e6 cells / mL. Nalm6 cells were cultured for up to ∼20 passages before discarding cultures are returning to low passage number frozen aliquots of initial cell line stock acquired from ATCC.

### In Vitro Proliferation Assays

After initial stimulation at a 1:4 E:T ratio with Nalm6 target cells eight days post non-viral editing, T cells were expanded in XVivo15 media + 5% FBS with 50 U/mL human IL-2 over the next two weeks. Every 2-3 days, T cells were counted by automated cell counter, and reseeded at concentrations of approximately 1e6 T cells / mL culture media. Total cumulative proliferation was calculated compared to the input number of CAR T cells for each of the four tested selection methods.

## DATA AVAILABILITY

All data supporting the findings of this study are available within the paper and its Supplementary Tables. An annotated list of all gRNA sequences used in the study is available in Supplementary Table 1. An annotated list of all DNA constructs used in the study, as well as full DNA sequences for all constructs, is available in Supplementary Table 2. Annotated genebank files for all DNA constructs used in the study are available upon request.

## ACKNOWLEDGEMENTS

We thank the members of the Satpathy lab for stimulating discussions. This work was supported by a Career Award for Medical Scientists from the Burroughs Wellcome Fund (A.T.S.), a Lloyd J. Old STAR Award from the Cancer Research Institute (A.T.S.), the Parker Institute for Cancer Immunotherapy (A.T.S), a CRISPR Cures for Cancer Award (A.T.S.), and the California Institute for Regenerative Medicine (DISC2–13212, A.T.S.).

## COMPETING INTERESTS

T.L.R. is a founder of Arsenal Biosciences. A.T.S. is a founder of Immunai, Cartography Biosciences, and Prox Biosciences, an advisor to Zafrens, Pallando Therapeutics, and Wing Venture Capital, and receives research funding from Merck Research Laboratories.

### Contributions

T.L.R. and A.T.S. conceptualized the study. T.L.R. and A.T.S. wrote and edited the manuscript and all authors reviewed and provided comments on the manuscript. T.L.R., J.L., A.M., and O.T.N. performed experiments and analyzed data. T.L.R. and A.T.S. guided experiments and data analysis.

